# Fully accessible fitness landscape of oncogene-negative lung adenocarcinoma

**DOI:** 10.1101/2023.01.30.526178

**Authors:** Maryam Yousefi, Laura Andrejka, Monte M. Winslow, Dmitri A. Petrov, Gábor Boross

**Author notes:** Corresponding authors: Monte M. Winslow, Stanford University School of Medicine | 279 Campus Drive, Beckman Center B256, Stanford, CA 94305. |, Dmitri Petrov, Stanford University Department of Biology | 327 Campus Drive, Bass Biology 232, Stanford, CA 94305 |, Gábor Boross, Stanford University Department of Biology | 327 Campus Drive, Bass Biology 230A, Stanford, CA 94305. |.

## Abstract

Cancer genomes are almost invariably complex with genomic alterations cooperating during each step of carcinogenesis. In cancers that lack a single dominant oncogene mutation, cooperation between the inactivation of multiple tumor suppressor genes can drive tumor initiation and growth. Here, we shed light on how the sequential acquisition of genomic alterations generates oncogene-negative lung tumors. We couple tumor barcoding with combinatorial and multiplexed somatic genome editing to characterize the fitness landscapes of three tumor suppressor genes NF1, RASA1, and PTEN, the inactivation of which jointly drives oncogene-negative lung adenocarcinoma initiation and growth. The fitness landscape was surprisingly accessible, with each additional mutation leading to growth advantage. Furthermore, the fitness landscapes remained fully accessible across backgrounds with additional tumor suppressor mutations. These results suggest that while predicting cancer evolution will be challenging, acquiring the multiple alterations required for the growth of oncogene-negative tumors can be facilitated by the lack of constraints on mutational order.

## INTRODUCTION

Fitness landscapes are a helpful conceptual aid in understanding evolutionary processes^1^. In a fitness landscape, genotypes are represented as nodes on a graph such that two genotypes are connected by an edge if they differ by a single allele and thus each genotype can be reached from the other by a single alteration^2^. Fitness values of each genotype are shown as an extra (height) dimension with adaptation represented as a hill-climbing adaptive walk. The key properties of evolutionary adaptation, such as its predictability and the ability of adaptation to reach the fittest genotype in a deterministic walk, are determined by the shape of the landscape, specifically by its ruggedness^3^. The ruggedness of the fitness landscape is defined by the frequency of sign epistasis, a phenomenon where a mutation is advantageous in one genetic context but disadvantageous in another. Sign epistasis determines whether an adaptive walk can end up stalled on a local fitness peak and whether only some combinations of mutations are sequentially adaptive. Intriguingly, many previous studies found that rugged fitness landscapes are common across diverse evolutionary systems^3–7^. This suggests that early adaptive mutations can strongly affect the direction of the adaptive process and limit the accessible adaptive paths.

Cancers are driven by the sequential acquisition of genetic driver mutations and thus are canonical examples of adaptive walks on a fitness landscape. Nonetheless, full fitness landscapes for cancer remain poorly understood due to the difficulty of inferring the fitness of growing tumors, the combinatorially large numbers of intermediate genotypes for even moderately genetically complex tumors, and the fact that tumors are generally analyzed once they have accumulated a large number of mutations with the order of the early steps hidden from the study. Despite the importance of defining the properties of cancer fitness landscapes for understanding tumorigenesis, we currently lack experimental systems to quantify the fitness of nascent tumors of diverse sequential genotypes growing *in vivo*. Defining cancer fitness landscapes could help to determine cancer progression and its therapeutic vulnerabilities. Human cancer genomics has uncovered genomic alterations that are mutually exclusive with one another, and we and others have found sign epistasis between oncogene and tumor suppressor mutations that suggests that cancer fitness landscapes might be rugged and not fully accessible^8–10^.

Oncogene-negative tumors represent ~20% of lung adenocarcinoma cases and affect ~150,000 patients each year worldwide^11–14^. These tumors lack oncogene mutations, and a large fraction of them could be driven by the combinatorial inactivation of tumor-suppressive genes^14,15^. We recently identified combinatorial inactivation of *Nf1, Rasa1*, and *Pten* and activation of MAPK and PI3K pathway as a potent driver of a subset of oncogene-negative lung tumors ^14^. While oncogene-negative tumors represent a large fraction of lung cancer, they likely require the coincident inactivation of multiple tumor suppressor genes to drive growth comparably to a single oncogene alteration. This raises the question of the extent to which genetic interaction among individual tumor suppressor mutations affect fitness during cancer evolution. Does inactivation of each gene confer a growth benefit regardless of genetic context, and hence form a fully accessible fitness landscape, or do they form a rugged fitness landscape where only certain orders of mutations are favored by selection?

Here, we characterized the full fitness landscape of autochthonous oncogene-negative lung tumors with combinatorial inactivation of the NF1, RASA1 and PTEN tumor suppressor genes. We used combinatorial and multiplexed somatic CRISPR/Cas9 genome editing and tumor barcoding to quantify the accessibility of the Nf1, Rasa1, Pten triple-mutant genotype via all possible single steps. We further investigated how the shape of the fitness landscape is affected by the inactivation of other tumor suppressor genes. We uncover an unexpectedly smooth landscape that remains accessible even in the presence of additional tumor suppressor alterations.

## RESULTS

### Quantitative analysis of the size and number of tumors of each stepwise genotype towards *Nf1;Rasa1;Pten* triple-mutant lung cancer

To quantify the impact of diverse genotypes on tumor initiation and growth, we developed a method based on tumor barcoding coupled with high-throughput barcode sequencing (Tuba-seq)^16–18^. Tuba-seq uses lentiviral-based DNA barcoding coupled with CRISPR-Cas9 genome editing and genetically engineered mouse alleles to generate and quantify the number and size of tumors of many different genotypes in parallel (**Supplementary Fig. 1a)**. Each lentiviral vector encodes Cre recombinase, a sgRNA, as well as a two-component barcode region with a fixed sgRNA identifier (sgID) that labels the vector and a random barcode (BC) that uniquely tags each clonal tumor that arises from the initial transduced cell.

We previously used Tuba-seq to identify combinations of tumor suppressor genes whose inactivation is capable of driving lung adenocarcinoma in the absence of oncogene mutations. Specifically, we discovered that coincident inactivation of *Nf1, Rasa1*, and *Pten* is a potent driver of oncogene-negative lung adenocarcinoma^14^. To investigate the contribution of single, double, and triple mutations of *Nf1, Rasa1*, and *Pten* to the genesis of oncogene-negative lung adenocarcinomas, we previously initiated tumors in *R26*^*LSL-Tomato*^*;H11*^*LSL-Cas9*^ (*TC*) and *Trp53*^*f/f*^*;TC* mice with a pool of eight uniquely dual barcoded lentiviral vectors with sgRNAs targeting *Nf1, Rasa1*, and *Pten* (**Fig. 1a)**. This pool contains lentiviral triple sgRNA vectors that target each gene alone, in pairwise combinations, and all three together (Lenti-sg*TS*^*Triple-pool*^*/Cre*, **Fig. 1a**)^14^. These mice developed thousands of clonal tumors of different genotypes (**Supplementary Fig. 1b-c**). Our previous Tuba-seq analysis indicated that most of the tumor burden in *TC* and *Trp53*^*f/f*^*;TC* mice was from tumors with concomitant inactivation of all three tumor suppressor genes^14^. However, we had not previously used our high-resolution tumor barcoding data to build a fitness landscape to understand the potential evolutionary paths to the triple mutant state, nor had we compared the tumorigenic potential between the p53-proficient and -deficient backgrounds.

**Figure 1.**
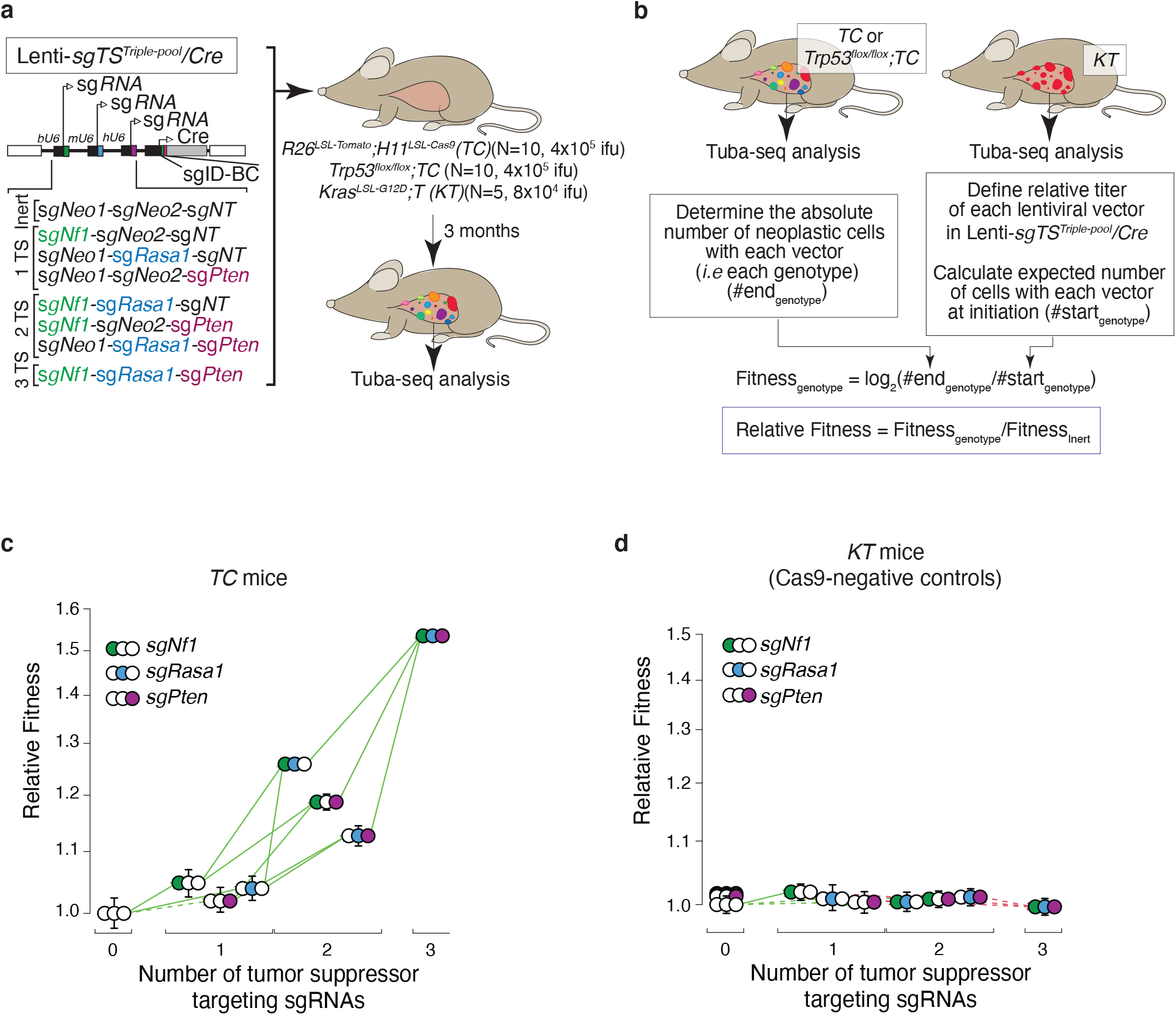
The fitness landscape of *Nf1;Rasa1;Pten* oncogene-negative lung adenocarcinoma is fully accessible. **a**, Schematic of barcoded triple sgRNA vectors in the Lenti-sg*TS*^*Triple-pool*^*/Cre* pool for CRISPR/Cas9-mediated inactivation of all combinations of *Nf1, Rasa1*, and *Pten* in *R26*^*LSL-*^ _*Tomato*_*;H11*^*LSL-Cas9*^ (*TC*), *Trp53*^*flox/flox*^*;TC* mice, and Cas9-negative *Kras*^*LSL-G12D*^*;T* mice. sgNeo1, sgNeo2, and sgNT are inert sgRNAs. The number and size of tumors initiated with each vector can be quantified by tumor barcoding coupled with high-throughput sgID-BC sequencing (Tuba-seq). Mouse genotype, mouse number, and viral titer (ifu; infectious units) are indicated. Tuba-seq was performed on bulk tumor-bearing lungs 3 months after tumor initiation. Data was generated in Yousefi, Boross *et al*., Cancer Research 2022. **b**, Overview of our method to calculate Relative Fitness of each cancer genotype (Methods). **c-d**, Fitness landscape for *TC* (**c**) and *Kras*^*LSL-G12D*^*;T* (Cas9-negative control)(**d**) mice. Fitness for tumors of each single and double mutant genotype, as well as those with all three tumor suppressor gene targeting sgRNAs are shown relative to the triple inert vector. Green arrows indicate increased fitness, red arrows indicate reduced fitness, solid line indicates significance (p-value < 0.05). Whiskers show 95% confidence intervals from bootstrap resampling of tumors.

### The fitness landscape of *Nf1;Rasa1;Pten* triple-deficient lung cancer is completely accessible

Generating Tuba-seq data on tumors initiated with lentiviral vectors that create all possible combinatorial mutations of *Nf1, Rasa1*, and *Pten* should enable the generation of a complete fitness landscape across all genotypes. We approximated the relative fitness of each genotype by calculating the growth rate using data on tumor number and size (**Fig. 1b** and **Methods**). Across these three tumor suppressor genes there are six possible paths on the landscape, *i*.*e* six possible routes from the genotype with the lowest fitness to the one with the highest fitness (**Supplementary Fig. 2a)**. As anticipated, we found that *Nf1;Rasa1;Pten* triple mutant tumors had the highest relative fitness in *TC* mice (**Fig. 1c**). Interestingly, all mutational routes leading to the *Nf1;Rasa1;Pten* triple mutant genotype were accessible, with each additional mutation increasing fitness beyond the fitness of the less complex genotypes of origin (**Fig. 1c**). These results are consistent with a model in which inactivation of each tumor suppressor gene is advantageous on all backgrounds regardless of whether the other two tumor suppressor genes are wild-type or mutant.

To assess the robustness of our relative fitness estimates, we performed a series of additional experiments. As an initial negative control for our method, we analyzed the tumors initiated with Lenti-sg*TS*^*Triple-pool*^*/Cre* in control *Kras*^*LSL-G12D*^*;T* (*KT*) mice, which lack Cas9 (**Fig. 1a**). In these mice, all sgRNAs are inert, and tumors with different sgIDs have the same genotype. As anticipated, our method uncovered a fitness landscape that was not accessible and that lacked the gains in fitness for tumors initiated with vectors with sgRNA targeting increasing numbers of tumor suppressor genes (**Fig. 1d**). Next, we performed a different method in which we used bootstrap resamples of mice as well as tumors, which had minimal impact on the overall significance of the fitness differences across the landscape in *TC* mice (**Supplementary Fig. 2b**).

In lung cancer models using lentiviral vectors, a small percent of tumors arise from cells that were transduced by more than one vector^14,16^. In the context of our experiments to map fitness, these multiple transduction events could impact tumor growth (except for tumors with the *Nf1;Rasa1;Pten* triple vector as they have the maximum numbers of possible mutations). We know that multiple infections must occur as we observe a small number of tumors containing the triple sgInert vector. These tumors are certainly the result of this vector co-transducing an initial cell with a vector that encodes one or more tumor suppressor gene targeting sgRNAs. We performed a series of analyses taking into account the expected rates of multiple transduction events and corrected our data to account for this effect. The number of sgInert tumors in *TC* mice was used to calculate the rate of coinfection in our experiments. We then used the rate of coinfection to remove a fraction of tumors of each genotype, picking the tumors to remove based on the size distribution of *Nf1*;*Rasa1*;*Pten* triple mutant size distribution (**Methods**). Importantly, the landscape remained accessible in the absence of correction for multiple transduction events (**Supplementary Fig. 2c**) as well as when we used an alternate method to correct for the impact of multiple transduction events (**Supplementary Fig. 2d**, see **Methods**). Our results show that the observed accessibility of the fitness landscape is robust to different methods to correct for multiple transduction events. These results are also consistent with experimental evidence from our previous study in which single and double mutant genotypes lead to some neoplastic growths *in vivo*^14^.

### *Trp53*-deficiency reduced tumor initiation without impacting the fully accessible landscape

In oncogenic KRAS, EGFR, and BRAF-driven lung cancer models, *Trp53* inactivation increases tumor initiation/early tumor growth and tumor size^9,17,19^. Our data allowed us to test whether Trp53 inactivation has a similar effect on the growth of oncogene-negative tumors, especially in case of the potent *Nf1;Rasa1;Pten* triple-deficient tumors. Similarly to the effects in the oncogene-driven tumors, multiple metrics of tumor size indicated that all tumors, as well as the *Nf1;Rasa1;Pten* triple-deficient tumors were larger in *Trp53*^*f/f*^*;TC* mice relative to *TC* mice (**Fig. 2a-b,f** and **Supplementary Fig. 3a-d**,**f**). In contrast, *Trp53* inactivation reduced overall tumor number (*i*.*e* number of tumors estimated to have >50 neoplastic cells), as well as *Nf1;Rasa1;Pten* triple-deficient tumor number (**Fig. 2c-d,f** and **Supplementary Fig. 3f**). These results agree with our previous visual analysis of surface tumor number in these mice^14^ (**Fig. 2e** and **Supplementary Fig. 3e**). However, the reduction in tumor initiation is the opposite of the effect of *Trp53*-deficiency in oncogenic KRAS, BRAF and EGFR-driven lung cancer models^8,9,17^. Thus, *Trp53* inactivation has a uniquely divergent effect on tumor initiation and tumor size in oncogene-negative tumors.

**Figure 2.**
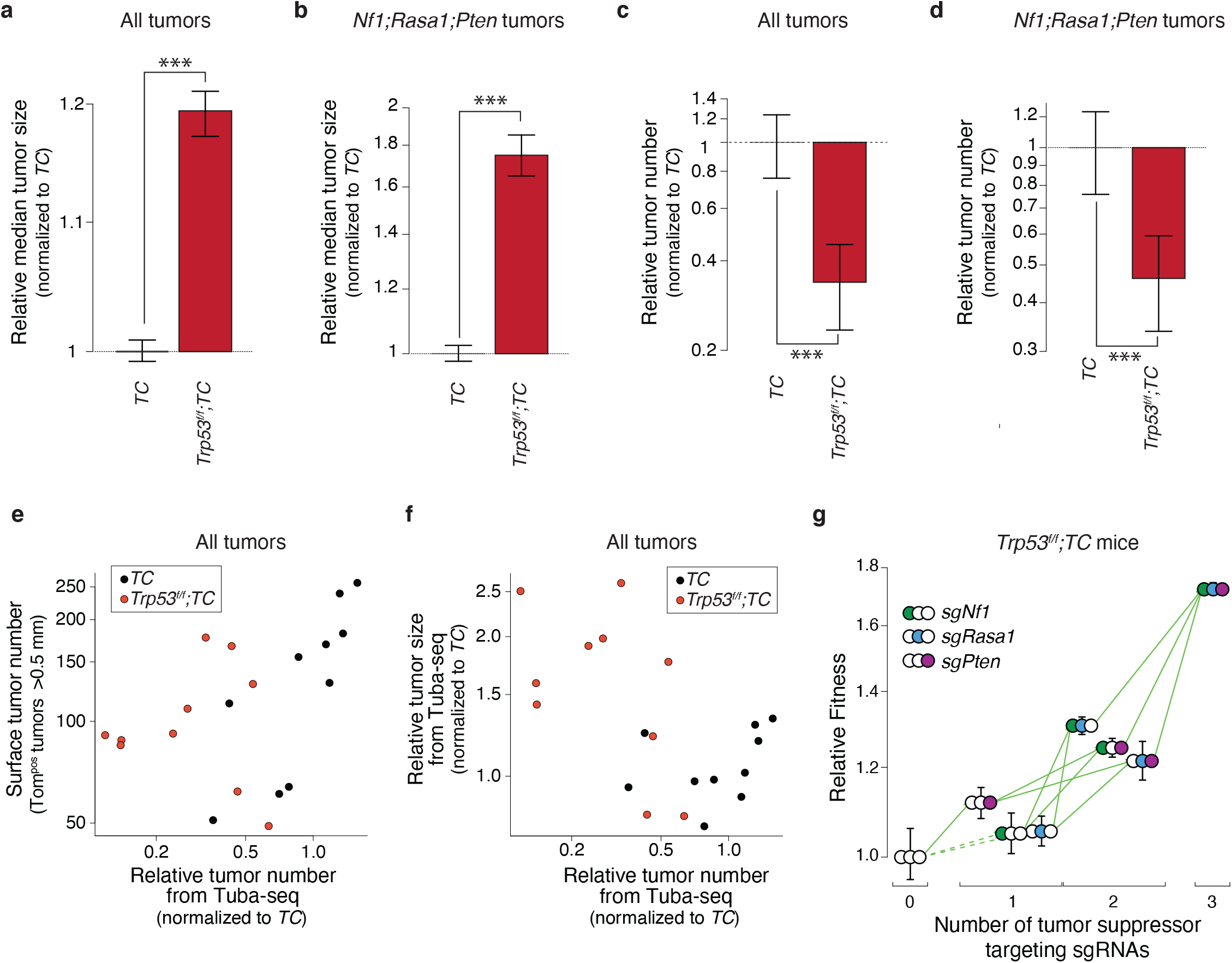
The fitness landscape of *Nf1;Rasa1;Pten* oncogene-negative lung adenocarcinoma remains fully accessible despite otherwise strong impacts of Trp53-deficiency. **a-b**, Effect of *Trp53* inactivation on the median size of all tumors (**a**) and of *Nf1;Rasa1;Pten* tumors (**b**). Effects are shown relative to *TC* mice. Whiskers show 95% confidence intervals from bootstrap resampling. ***, P < 0.001 based on a bootstrap resampling. **c-d**, Effect of *Trp53* inactivation on the number of all tumors (**c**) and of *Nf1;Rasa1;Pten* tumors (**d**). Effects are shown relative to *TC* mice. Whiskers show 95% confidence intervals from bootstrap resampling. ***, P < 0.001 based on a bootstrap resampling. **e**, Comparison of Surface tumor number (from direct counting of Tomato^positive^ tumors >0.5 mm in diameter) with the Relative tumor number from Tuba-seq (the number of clonal exapansions with >50 neoplastic cells relative to the median of *TC* values). Each dot represents a mouse. Data from all tumors is shown. **f**, Comparison of the Relative tumor size determined from Tuba-seq (the median number of neoplastic cells relative to the median of *TC* values) with the Relative tumor number from Tuba-seq (the number of clonal expansions with >50 neoplastic cells relative to the median of *TC* values). Each dot represents a mouse. Data from all tumors is shown. **g**, Fitness landscape for *Trp53*^*f/f*^*;TC* mice. Fitness for tumors of each single and double mutant genotype, as well as those with all three tumor suppressor targeting sgRNAs are shown relative to the triple inert vector. Green arrows indicate increased fitness, red arrows indicate reduced fitness, solid line indicates significance (p-value < 0.05). Whiskers show 95% confidence intervals from bootstrap resampling of tumors.

Next, we mapped the fitness landscape of *Nf1, Rasa1*, and *Pten* mutations on the background of *Trp53* deficiency in *Trp53*^*f/f*^*;TC* mice. Similar to our observations in *Trp53* proficient *TC* mice, the *Nf1;Rasa1;Pten* triple mutant genotype had the highest relative fitness. Importantly, despite the overall effects of *Trp53* deficiency on tumor number and size, the landscape remained fully accessible (**Fig. 2g** and **Supplementary Fig. 4a-c**).

### Quantification of the overall tumorigenicity of the *Nf1*;*Rasa1*;*Pten* triple-mutant genotype across *Lkb1* and *Keap1* deficient backgrounds

While the fitness landscape of *Nf1*;*Rasa1*;*Pten* lung tumors was fully accessible in *Trp53*-proficient and -deficient backgrounds, changes in the genetic background often influence epistatic interactions, and thus change landscape accessibility. Therefore, we next quantified whether the effects of combinatorial *Nf1, Rasa1*, and *Pten* inactivation change on other genetic backgrounds. We chose loss-of-function mutations in *Keap1* and *Lkb1*(*Stk11*) which are known tumor suppressor genes that are frequently mutated in human lung cancer and have been shown to suppress growth in models of oncogenic KRAS-driven lung adenocarcinoma^16,17,20–22^. We initiated tumors with Lenti-sg*TS*^*Triple-pool*^*/Cre* in *TC, Keap1*^*f/f*^*;TC, Lkb1*^*f/f*^*;TC* and control *Kras*^*LSL-G12D*^*;T* mice (**Fig. 3a**). After 3 months of tumor growth, we first analyzed tumor burden by direct fluorescent imaging and histology. Unexpectedly, *Keap1*^*f/f*^*;TC* mice developed many fewer tumors than *TC* mice as assessed by direct fluorescence and histology (**Fig, 3b-c**). Conversely, *Lkb1*^*f/f*^*;TC* mice visually had many tumors that appeared somewhat larger than those in *TC* mice (**Fig. 3b-c**).

**Figure 3.**
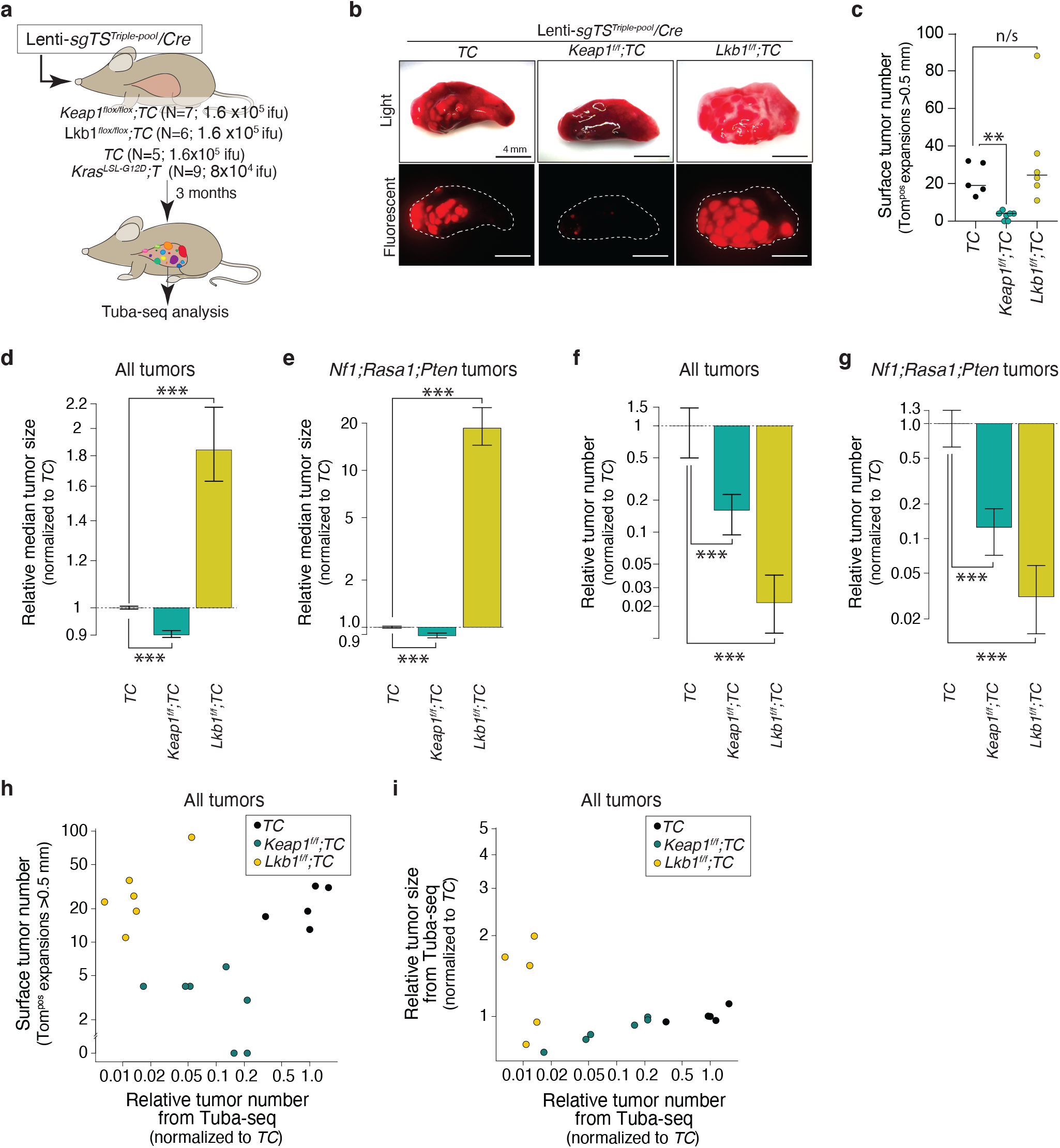
Inactivaton of the Keap1 or Lkb1 “tumor suppressors” reduces *Nf1;Rasa1;Pten* oncogene-negative tumor initiation with divergent effects on tumor size. **a**, Schematic of experiments to assess the effect of *Keap1* and *Lkb1* inactivating background mutations on combinations of *Nf1;Rasa1;Pten* inactivation. Mouse genotype, mouse number, and viral titer (ifu; infectious units) are indicated. Tuba-seq was performed on bulk tumor-bearing lungs 3 months after tumor initiation. **b**, Representative light and fluorescence images of lung lobes from the indicated genotypes of mice. Lung lobes are outlined with white dotted lines. Scale bars = 4 mm **c**, The number of tumors (Tomato^positive^ tumors >0.5mm in diameter) quantified by direct counting. Each dot represents a mouse, and the bar is the median. The genotypes of the mice are indicated. **, P < 0.01; and n/s, P > 0.05 (not significant) based on a bootstrap resampling. **d-e**, Effect of *Keap1* or *Lkb1* inactivation on the median size of all tumors (**d**) and of *Nf1;Rasa1;Pten* tumors (**e**). Effects are shown relative to *TC* mice. Whiskers show 95% confidence intervals from bootstrap resampling. ***, P < 0.001 based on a bootstrap resampling. **f-g**, Effect of *Keap1* or *Lkb1* inactivation on the number of all tumors (**f**) and of *Nf1;Rasa1;Pten* tumors (**g**). Effects are shown relative to *TC* mice. Whiskers show 95% confidence intervals from bootstrap resampling. ***, P < 0.001 based on a bootstrap resampling. **h**, Comparison of Surface tumor number (from direct counting of Tomato^positive^ tumors >0.5mm in diameter) with the Relative tumor number from Tuba-seq (the number of clonal expansions with >50 neoplastic cells relative to the median of *TC* values). Each dot represents a mouse. Data from all tumors is shown. **i**, Comparison of the Relative tumor size determined from Tuba-seq (the median number of neoplastic cells relative to the median of *TC* values) with the Relative tumor number from Tuba-seq (the number of clonal expansions with >50 neoplastic cells relative to the median of *TC* values). Each dot represents a mouse. Data from all tumors is shown.

Next, we performed Tuba-seq on tumor-bearing lungs and quantified the effect of *Keap1* and *Lkb1* deficiency on overall tumor initiation and size. *Keap1* inactivation decreased overall tumor number by ~80%, while modestly reducing tumor size (**Fig. 3d-g** and **Supplementary Fig. 5b-c**). The impact of *Keap1* inactivation on tumor number and size was consistent across all tumors, as well as when we assessed only *Nf1;Rasa1;Pten* triple-deficient tumors (**Fig. 3d-g**). Thus, *Keap1* inactivation has an overall negative effect on tumorigenesis driven by inactivation of *Nf1, Rasa1*, and/or *Pten* with the greatest impact on tumor initiation and/or very early tumor growth.

The impact of *Lkb1* inactivation was even more interesting. *Lkb1* inactivation decreased overall tumor number by >95% (**Fig. 3f-g**), which seemed at odds with the gross examination of the tumor-bearing lung. However, the small number of tumors that did form were much larger than those in *TC* mice, with *Nf1;Rasa1;Pten* triple-mutant tumors in *Lkb1;TC* having >15-fold greater median tumor size than in *TC* mice (**Fig. 3d-e**). This dramatic increase in the size reconciles the visual observation of similar numbers of tumors in *Lkb1;TC* and *TC* mice, while quantitative analysis by Tuba-seq data indicated that *Lkb1;TC* mice had many fewer tumors (**Fig. 3h-i** and **Supplementary Fig. 5d-e**).

### Landscape accessibility is largely robust to changes in the genetic background

As the *Lkb1* and *Keap1* deficient backgrounds had substantial effects on overall oncogene-negative lung tumor growth in our experimental model, we next characterized their impacts on the fitness landscapes. Due to the very low number of tumors for less complex (inert, single and double mutant) genotypes on the *Lkb1*^*f/f*^*;TC* background, we could not construct a fitness landscape (**Supplementary Fig. 6a-b**). However, our method allowed us to generate fitness landscapes for the tumors in *TC, Keap1;TC*, and control *KT* mice. The landscape in *TC* mice was similar to that generated from our initial cohort of *TC* mice and the landscape in the control *KT* Cas9-negative mice was again very flat (**Fig. 4a** and **Supplementary Fig. 7d**). Interestingly, in the *Keap1*-deficient background, the landscape remained fully accessible (**Fig. 4b**). These effects were consistent across our different analysis methods (**Supplementary Fig. 7a-c**). Thus, despite greatly reducing tumor initiation, the ability of each additional mutation to increase fitness on the path to the *Nf1;Rasa1;Pten* triple deficient state was maintained.

**Figure 4.**
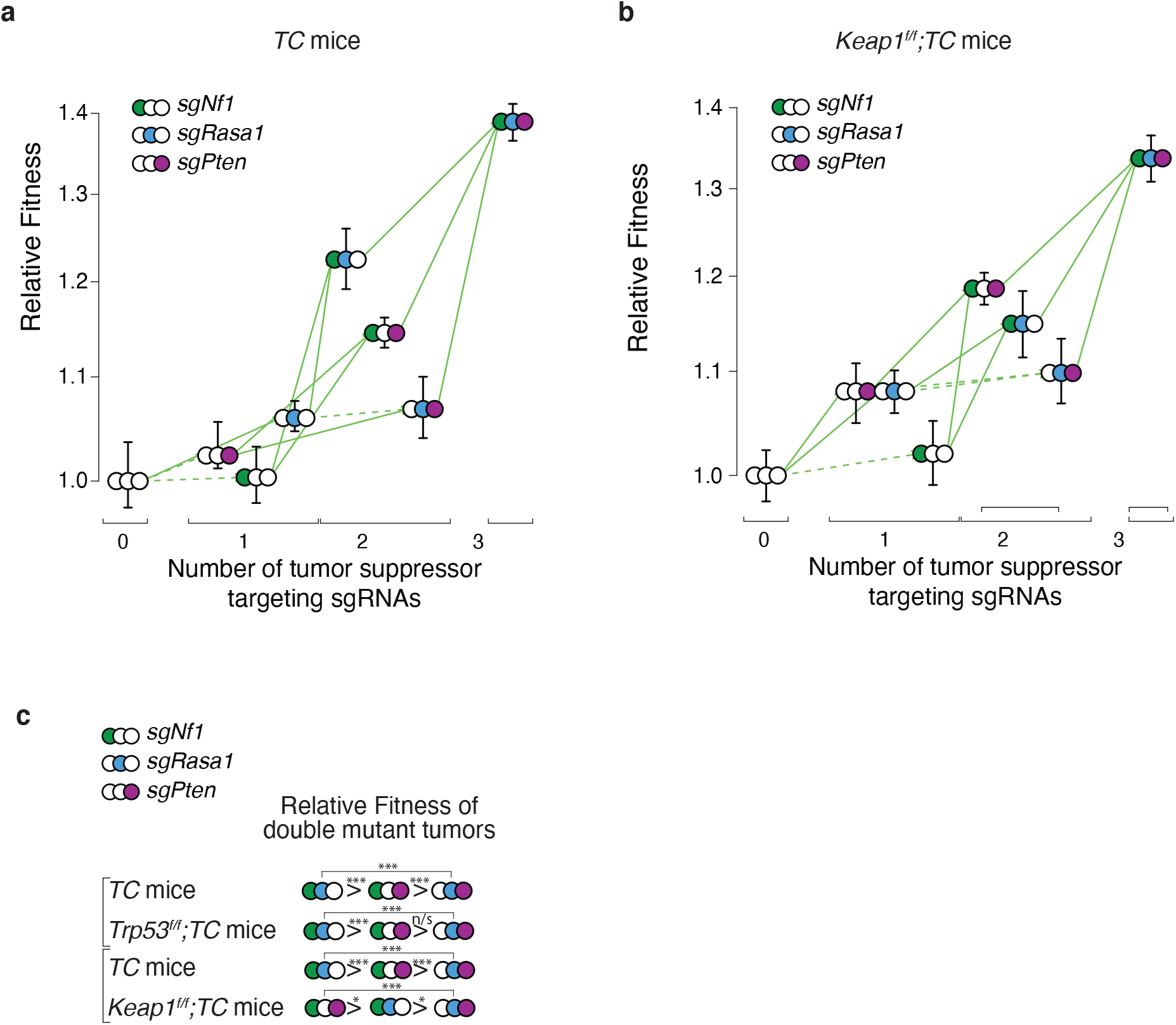
Keap1 inactivation changes the topology of the fitness landscape of *Nf1;Rasa1;Pten* oncogene-negative lung adenocarcinoma. **a-b**, Fitness landscape for *TC* (**a**) and *Keap1*^*flox/flox*^*;TC* (**b**) mice. Fitness for tumors of each single and double mutant genotype, as well as those with all three tumor suppressor gene targeting sgRNAs are shown relative to the triple inert vector. Green arrows indicate increased fitness, red arrows indicate reduced fitness, solid line indicates significance (p-value < 0.05). Whiskers show 95% confidence intervals from bootstrap resampling of tumors. **c**, Relative fitness of the double mutant genotypes from the *TC* and *Trp53*^*f/f*^*;TC* mice from Figure 1/2 and from *TC* and *Keap1*^*f/f*^*;TC* mice from Figure 4. *, P < 0.05; ***, P < 0.001; and n/s, P > 0.05 (not significant), based on a bootstrap resampling.

Despite the overall accessibility of the fitness landscape in both *Keap1*-proficient and -deficient backgrounds, the impact of additional mutation was not identical. In particular, the relative fitness of tumors of the different double mutant genotypes was different, with *Nf1;Rasa1* being most fit in the *TC* and *Trp53;TC* background and *Nf1;Pten* being the most fit in the *Keap1;TC* background (**Fig. 4c**). Thus, changes in the genetic background did impact the topology of the landscape, but it remained fully accessible.

## DISCUSSION

Our results suggest that the combinatorial inactivation of *Nf1, Rasa1*, and *Pten* forms a fully accessible fitness landscape. Thus, any of the six evolutionary routes is possible, with the inactivation of each gene increasing fitness beyond the preceding genotype. The overall lack of sign epistasis makes predicting cancer evolution challenging. On the other hand, while oncogene-negative tumors require the inactivation of multiple tumor suppressor genes to substitute for the lack of a strong oncogene alteration, gaining fitness advantage from each additional alteration increases the probability of acquiring all of the required mutations. Human genomics data suggest that oncogene-negative lung adenocarcinoma with high MAPK and PI3K pathway activation are likely generated from combinatorial mutations in diverse components of these pathways further expanding the available paths to the tumorigenic state^14^.

Contrary to the fitness landscapes we mapped in this study, the landscapes of many other evolutionary systems are ruggedand exhibit sign epistasis between mutations^3–7^. Mutations often have pleiotropic effects, with some effects being advantageous and other effects being detrimental. In one genetic context the beneficial effects can outweigh the detrimental effects, while in a different genetic context, this can be reversed. This leads to sign epistasis and a rugged fitness landscape. Why is the oncogene-negative fitness landscape smooth and thus fully accessible? We speculate that this could be because somatic cells and neoplastic cells *in vivo* are far from the highest possible fitness and individual tumor suppressor gene mutations have relatively weak effects. The combined effect could create sufficiently linear behavior of all the individual tumor suppressor mutations without encountering any significant curvature of the fitness function. This is often not the case for the large effect mutations observed in experimental evolution of microorganisms or in the tumor suppressor effects on the oncogene-driven tumors.

While the fitness landscape of combinatorial *Nf1, Rasa1*, and *Pten* inactivation remained fully accessible and robust to inactivation of *Trp53, Keap1* or *Lkb1*, those alterations dramatically changed overall tumor initiation and growth. Comparing the effects of *Trp53, Keap1* and *Lkb1* inactivation on *Nf1;Rasa1;Pten* mutant tumors with their impact on various oncogene-driven tumors underscore the context-dependency of these effects^8,9,17^. *Trp53* inactivation increased tumor size but had a negative effect on the initiation of oncogene-negative tumors. This observation was in contrast to the universally positive effect of *Trp53* deficiency on multiple models of oncogene-driven lung cancer^8,9,17,19^. Interestingly, while LKB1 and KEAP1 are generally regarded as tumor suppressors, several lines of evidence suggest that their role in cancer is more complex and context dependent^24^. *Lkb1* and *Keap1* inactivation has been shown to have detrimental effects on *in vivo* lung tumor initiation and growth across different genetic contexts^8,9,25,26^. Thus, while the detrimental effect of *Lkb1* and *Keap1* inactivation on initiation/early growth and the positive effect of *Lkb1* inactivation on the growth of oncogene-negative tumors could not have been predicted, it is consistent with these “tumor suppressors” having highly context-dependent effects.

Human tumors likely evolve through states in which tumor suppressor genes are heterozygously inactivated. However, as CRISPR/Cas9-genome editing generally generates homozygous gene inactivation, we could only study homozygous mutations in tumor suppressor genes. In the future, deciphering where the multiple middle steps of heterozygous mutants fall within cancer fitness landscapes would improve the resolution. The effects of heterozygous genotypes are likely to fall within the range of their homozygous counterpart and therefore are unlikely to render the fitness landscape inaccessible.

Most human tumors have complex genotypes, often including multiple inactivating alterations. Modeling complex cancer genotypes in genetically engineered mouse models has been greatly facilitated by somatic CRISPR/Cas9 genome editing, however broad-scale quantitative analysis of complex cancer genotypes remains limited^14,19^. Generating defined combinatorial alterations *in vivo* allows not only the analysis of tumorigenesis driven by these complex genotypes, but also the assessment of contribution of each stepwise genomic alteration to tumor fitness and evolution. Quantitative and comprehensive fitness maps of other genotypes (within and beyond lung cancer) could uncover general rules of how driver alterations interact and affect cancer evolution by creating rugged and smooth cancer fitness landscapes. While libraries of vectors with pairs of sgRNAs have been broadly employed^27–31^ generating large pools of vectors to inactivate greater numbers of genes while maintaining the ability to perform highly-quantitative clonal-level analysis will require additional methods. In the future, the analysis of tumors with a comprehensive combination of many other genotypes coupled with molecular profiling and external interventions like therapies should illuminate the underlying biology of adaptation and uncover exploitable vulnerabilities.

## METHODS

### Animal Studies

The use of mice for the current study has been approved by the Institutional Animal Care and Use Committee at Stanford University, protocol number 26696. *Kras*^*LSL-G12D/+*^ (Jax # 008179 (*K*)), *R26*^*LSL-tdTomato*^(*ai9*) (Jax # 007909 (*T*)), and *H11*^*LSL-Cas9*^ (Jax # 026816), *Keap1*^*flox*^, *Lkb1* ^*flox*^ (Jax # 014143), and *Trp53*^*flox*^ (Jax # 008462) mice have been previously described^32–38^. All mice were on a C57BL/6:129 mixed background. Tumors were initiated by intratracheal delivery of Lenti-Triple-sgRNA*/Cre* vectors. Mice were allowed to develop tumors for 3 months after viral delivery.

### Tumor barcode sequencing and analysis

Tuba-seq libraries were generated as described previously^14^. Briefly, genomic DNA was isolated from bulk tumor-bearing lung tissue followed by PCR amplification of the sgID-BC region from 32 μg of bulk lung genomic DNA using Q5 Ultra II High-Fidelity 2x Master Mix (New England Biolabs, M0494X). Unique dual-indexed primers were used to amplify each sample followed by purification using Agencourt AMPure XP beads (Beckman Coulter, A63881). The libraries were pooled based on lung weights to ensure even reading depth and sequenced (read length 2×150bp) on the Illumina HiSeq 2500 or NextSeq 500 platform (Admera Health Biopharma Services). Tuba-seq analysis of tumor barcode reads was performed as previously described (*37, 39*).

### Histology and immunohistochemistry

Lung lobes were inflated with 4% formalin and fixed for 24 hours, stored in 70% ethanol, paraffin-embedded, and 4 μm thick sections were used for Hematoxylin and Eosin (H&E) staining and immunohistochemistry (IHC). Anti-RFP (Rockland, 600-401-379), anti-TTF1(Abcam, ab76013), anti-UCHL1(Sigma, HPA005993), anti-TP63 (Cell Signaling Technology, 13109) IHC was performed using Avidin/Biotin Blocking Kit (Vector Laboratories, SP-2001), Avidin-Biotin Complex kit (Vector Laboratories, PK-4001), and DAB Peroxidase Substrate Kit (Vector Laboratories, SK-4100) following standard protocols.

### Data availability

Tuba-seq barcode sequencing data from the experiment in Figure 1 and 2 has been previously deposited in NCBI’s Gene Expression Omnibus (https://www.ncbi.nlm.nih.gov/geo/) (GSE174393). Tuba-seq barcode sequencing data from the experiment in Figure 3 and 4 has been deposited in NCBI’s Gene Expression Omnibus (GSE223678).

### Calculation of relative fitness of each genotype of tumors

Fitness of each tumor genotype was approximated by the growth rate of tumors. Fitness was calculated for each tumor genotype in a given mouse strain by considering the expected number of cells with each lentiviral vector at the start of the experiment (#start_*genotype_mouse-strain*_) and number of neoplastic cells with each lentiviral vector after tumor growth at the end of the experiment (#end_*genotype_mouse-strain*_).

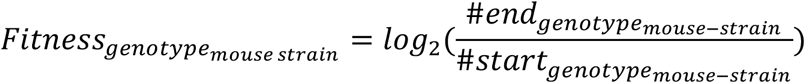

#end_*genotype_mouse-strain*_ is the sum of all neoplastic cells in all tumors of that genotype at the end of the experiment from all mice of a given strain (for example *TC* mice in Figure 1).

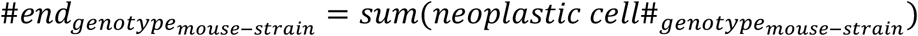

#start_*genotype_mouse-strain*_ is the expected number of cells of that genotype at the start of the experiment. #start_*genotype_mouse-strain*_ was determined from the number of tumors with each viral vector (titer_*genotype*_) from *KT* Cas9-negative control mice. In *KT* mice, the number of tumors with each vector (number of unique BCs associated with each sgID) represents the exact titer of each vector (titer_genotype_) in the Lenti-sg*TS*^*Triple-pool*^*/Cre* pool. To put #start_genotype_ in the context of the most potent genotype in the given mouse strain, we normalized the number of tumors with each lentiviral vector in *KT* mice (titer_genotype_) to the number of *Nf1;Rasa1;Pten* triple mutant tumors in *KT* mice (titer_*Nf1;Rasa1;Pten*_) and then multiplied this by the total number of *Nf1;Rasa1;Pten* triple mutant tumors across all mice of a given strain (TumorNumber_*Nf1;Rasa1;Pten_mouse-strain*_). Thus, #start_*genotype_mouse-strain*_ represents the titer-corrected expected number of tumors of that genotype if that genotype was as potent as triple *Nf1:Rasa1;Pten* mutation in that strain. As an example, #start for single mutant genotype *Nf1* in *TC* mice (#start _*Nf1_TC*_) was calculated as;

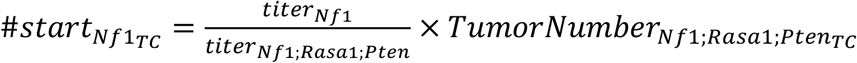

Finally, we calculated “Relative Fitness” within each mouse strain by normalizing the fitness of each genotype to the fitness of tumors with the lentiviral vector with three Inert sgRNAs (Lenti-sgNeo1-sgNeo2-sgNT/Cre; inert tumors).

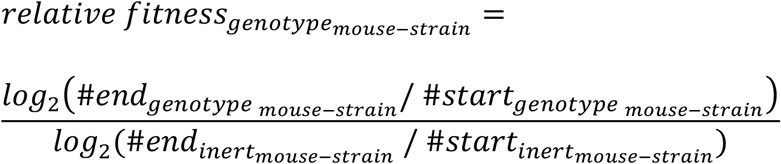

### Methods to control for multiple transductions

The most potent genotype in our experiments was the *Nf1;Rasa1;Pten* triple mutant generated by transduction with the Lenti-sg*Nf1*-sg*Rasa1*-sg*Pten*/Cre vector. We found that in our *in vivo* experiments that a small percent of tumors arise from lung epithelial cells that were initially transduced by more than one lentiviral vector^14^. We refer to these as multiple transductions. Multiple transductions influence our ability to estimate fitness as a subset of tumors identified as a given genotype (e.g. *Nf1* single mutant tumors which arise from a cell transduced with the Lenti-sg*Nf1*/Cre vector) are in fact double mutants or even *Nf1;Rasa1;Pten* triple mutant due to multiple transduction of the initial cells with a vector with complementary sgRNAs (e.g. with the Lenti-sgNf1-sg*Rasa1-sgPten*/Cre vector). To quantify the fraction of tumors that arise from multiple transductions (multiple transduction rate), we calculated the ratio of the number of tumors with greater than a given number of neoplastic cells (minimum cell number cutoff, X) for a given genotype versus the number of tumors with greater than the same minimum cell number cutoff for triple mutants:

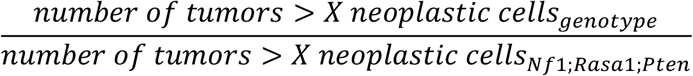

As we increase the minimum cell number cutoff (X), the ratio will decrease and asymptotically approach the true multiple transduction rate. The reason behind this is that as we increase the minimum cell number cutoff, the proportion of tumors co-transduced with the most potent triple mutant genotype among all our tumors increases, as the larger the tumor, the less likely it is to be from a genotype other than the *Nf1;Rasa1;Pten* triple mutant. Therefore, the minimum ratio across all minimum cell number cutoff (X) values was used as the multiple transduction rate. The multiple transduction rate provides us the expected number of tumors (n) for any given genotype that are the result of multiple transductions. Next, to control for the effect of multiples transduction, we used the distribution of *Nf1;Rasa1;Pten* triple mutant tumors to remove the n tumors from the data for each other genotype. For the n tumors to remove, we took the n-quantiles of the *Nf1;Rasa1;Pten* triple mutant tumor size distribution. For each of these quantile values, we chose the tumor of the given genotype that was closest to the quantile in size and remove that tumor from the dataset for the given genotype. We used the sgInert multiple transduction rate in *TC* mice for all genotypes and all mouse strains in all analysis and figures except **Figure 2e-f, 3h-i** and **Supplementary Figure 1b-c, 2c-d, 3e-f, 4a-b, 5d-e, 6a-b** and **7a-b**. This is logical, as multiple transductions should not be influenced by the sgIDs, sgRNA sequences, or mouse genotypes. Furthermore, we do not expect growth of tumors with Lenti-sgInert/vectors in *TC* mice, unless those initial cells have multiple transductions that occur with vectors with tumor suppressor gene targeting lentiviral vectors.

As an alternative method, we also quantified the multiple transduction rate for all genotypes of tumors separately and used that for multiple transduction correction (multiple transduction correction Method #2, see **Supplementary Fig. 2c, Supplementary Fig. 4b, Supplementary Fig. 7b**). The results and conclusions were unchanged when we used this alternate multiple transduction correction method.

### Statistical analysis

All statistical analyses were performed using the R software environment. For all bar plots showing relative tumor size and tumor number and for fitness landscape plots, p-values and 95% confidence intervals (represented by whiskers) were calculated using bootstrap resampling (10,000 repetitions). Bootstrapping was done by random resampling with replacement all the tumors in all mice of a given strain. When indicated in the figure legend and in cases of barplots showing tumor number, a nested bootstrap approach was used where first all the mice from a given strain were resampled with replacement and then all tumors in this virtual mouse cohort were resampled again with replacement.

## ACKNOWLEDGEMENTS

We thank the Stanford Veterinary Animal Care Staff for expert animal care; Leo Chen and Min K. Tsai for technical assistant; Chuan Li, Paloma Ruiz and other members of the Winslow and Petrov laboratories for helpful discussions and reviewing the manuscript. M. Yousefi was supported by a Stanford University School of Medicine Dean’s fellowship, an American Lung Association grant, and an NIH Ruth L. Kirschstein National Research Service Award (F32-CA236311). G. Boross was supported by Tobacco-Related Disease Research Program Postdoctoral Fellowships (T31FT-1619). This work was supported by NIH R01-CA231253 (to M.M. Winslow and D.A. Petrov), NIH R01-CA230919 (to M.M. Winslow) and NIH R01-CA234349 (to M.M. Winslow and D.A. Petrov), as well as by the Stanford Cancer Institute, an NCI-designated Comprehensive Cancer Center.

## Notes

### Competing Interest Statement

D.A.P. and M.M.W. are founders of, and hold equity in, D2G Oncology Inc.

### Summary of Updates

Mistakes in text and figures corrected.

