## Supplementary Figures 1-7. for "Fully accessible fitness landscape of oncogene-negative lung adenocarcinoma"

**a**

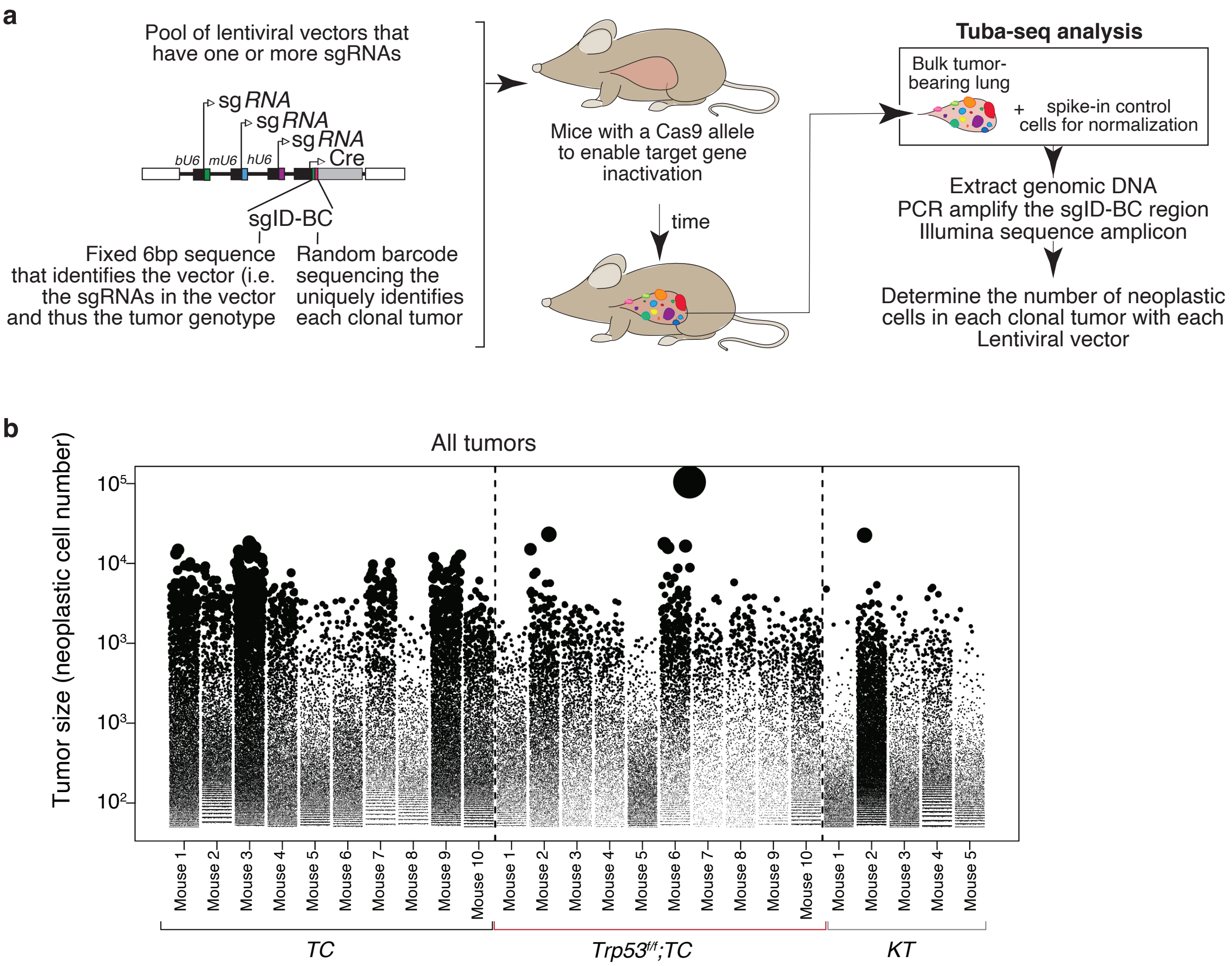


**c**

| | *TC* | *Trp53$^{f/f}$;TC* | *KT* |
|---|---|---|---|
| *Nf1;Rasa1;Pten* | 2438 | 1126 | 1836 |
| *Nf1;Rasa1* | 1317 | 525 | 1486 |
| *Nf1;Pten* | 1926 | 562 | 1887 |
| *Rasa1;Pten* | 1244 | 392 | 1520 |
| *Nf1* | 959 | 254 | 1706 |
| *Rasa1* | 1458 | 408 | 2740 |
| *Pten* | 699 | 223 | 1024 |
| *Inert* | 868 | 237 | 1634 |

**Supplementary Figure 1. *TC* and *Trp53$^{f/f}$;TC* mice with Lenti-sgTS$^{triple\text{-}pool}$/Cre-induced tumors develop many clonal tumors**
**a,** Overview of tumor barcoding coupled with high-throughput barcode sequencing (Tuba-seq).
**b,** Gitter plot of tumor sizes in each mouse of each genotype. Each dot represents a tumor (>50 neoplastic cells). Sizes of dots scale with tumor size. Each column represents a mouse. All tumors generated with any of the lentiviral vectors are shown.
**c,** Average total number of clonal tumors with each lentiviral vector in each mouse of each strain. Tumor numbers are the sum of all tumor with the given lentiviral vector across all mice of each genotype, divided by the number of mice of that genotype. Tumors are defined as clonal expansions estimated to have >50 neoplastic cells. These numbers reflect the total number of barcoded tumors prior to removal of tumors to reduce the impact of multiple transduction events (see **Methods**).

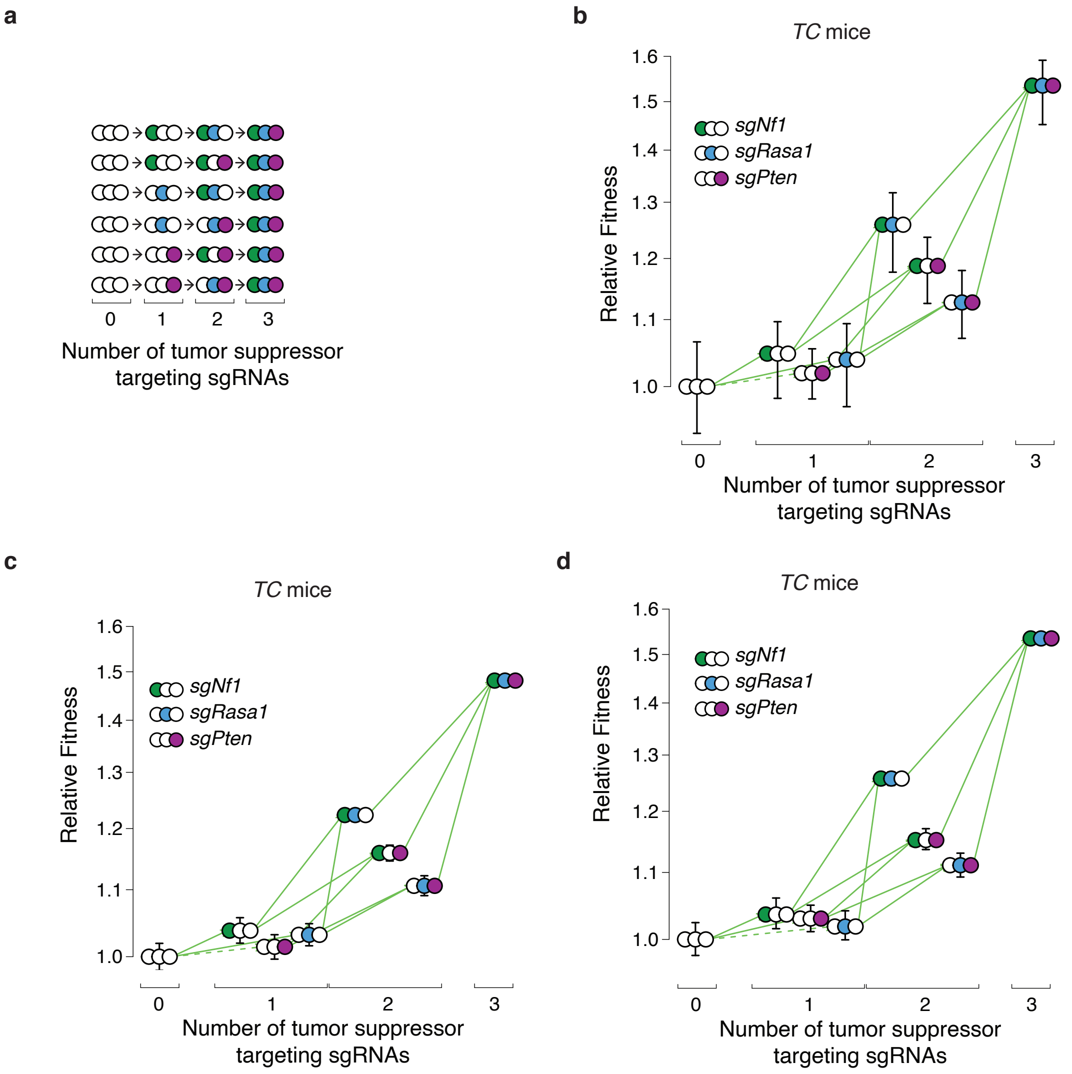


**Supplementary Figure 2. Estimates of fitness for tumors in *TC* mice are robust to different methods for multiple transduction correction**

**a,** There are six possible paths from wild type cells to the *Nf1;Rasa1;Pten* triple mutant genotype.

**b-d**, Fitness landscape of tumors in *TC* mice with multiple transduction correction as in Figure 1c but with 95% confidence intervals calculated through bootstrap resampling of both mice and tumors (**b**), without any multiple transduction corrrection (**c**), and with multiple transduction correction using method #2 (Methods)(**d**). Fitness for tumors of each single and double mutant genotype, as well as those with all three tumor suppressor targeting sgRNAs are shown relative to the triple inert vector. Green arrows indicate increased fitness, solid line indicates significance (p-value < 0.05). Whiskers in c and d show 95% confidence intervals from bootstrap resampling of tumors.

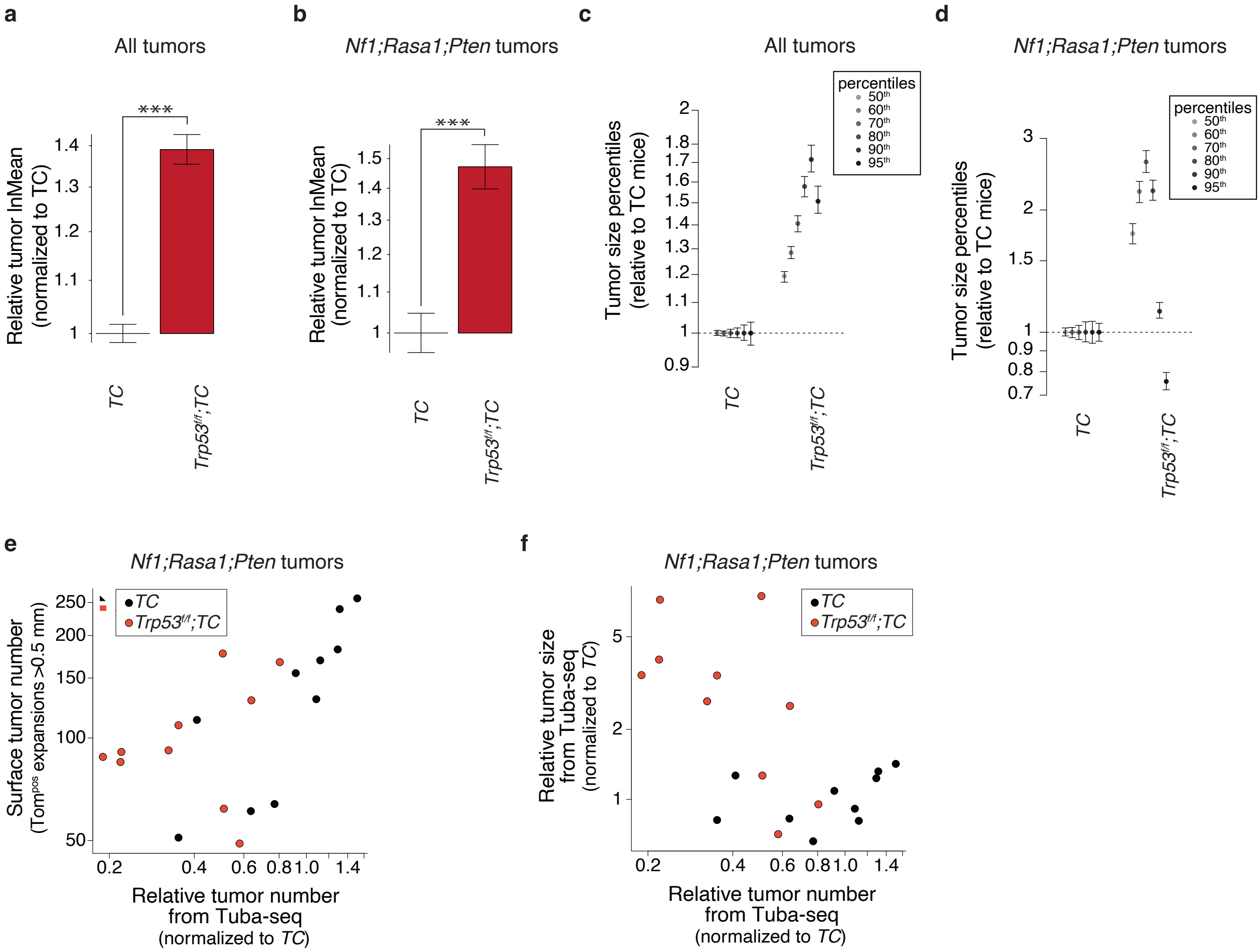


**Supplementary Figure 3. *Trp53* inactivation increases tumor size but reduce tumor initiation/early expansion**
**a-b,** Estimates of mean tumor size fitting a log-normal distribution (lnMean) are shown relative to the lnMean values in *TC* mice. Whiskers show 95% confidence intervals from bootstrap resampling. ***, P < 0.001 based on a bootstrap resampling.
**c-d,** Tumor sizes at indicated percentiles relative to the same percentile of tumor sizes in *TC* mice. Whiskers show 95% confidence intervals from bootstrap resampling.
**e,** Comparison of Surface tumor number (from direct counting of Tomato$^{positive}$ tumors >0.5mm in diameter) with the Relative tumor number from Tuba-seq (the number of clonal expansions with >50 neoplastic cells relative to the median of *TC* values). Each dot represents a mouse. Data from *Nf1;Rasa1;Pten* triple mutant tumors is shown.
**f,** Comparison of the Relative tumor size determined from Tuba-seq (the median number of neoplastic cells relative to the median of *TC* values) with the Relative tumor number from Tuba-seq (the number of clonal expansions with >50 neoplastic cells relative to the median of *TC* values). Each dot represents a mouse. Data from *Nf1;Rasa1;Pten* triple mutant tumors is shown.

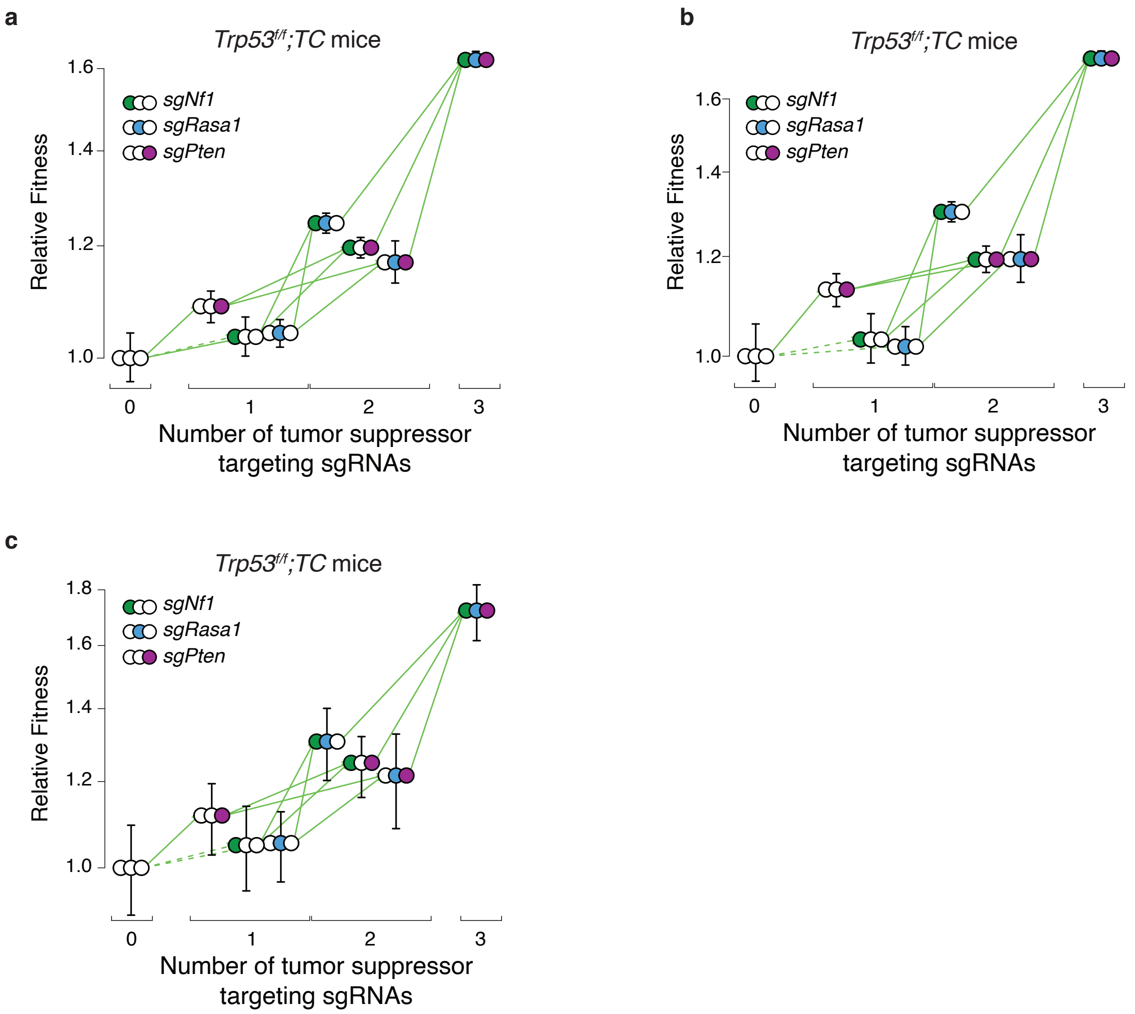


**Supplementary Figure 4. Estimates of fitness for tumors in *Trp53$^{f/f}$;TC* mice are robust to different methods for multiple transduction correction.**
**a-c,** Fitness landscape of tumors in *Trp53$^{f/f}$;TC* mice without any multiple transduction corrrection (**a**), with multiple transduction correction using method #2 (Methods)(**b**), and with multiple transduction correction as in Figure 2g but with 95% confidence intervals calculated through bootstrap resampling of both mice and tumors (**c**). Fitness for tumors of each single and double mutant genotype as well as those with all three genes inactivated are shown relative to inert. Green arrows indicate increased fitness, solid line indicates significance (p-value < 0.05). Whiskers show 95% confidence intervals.

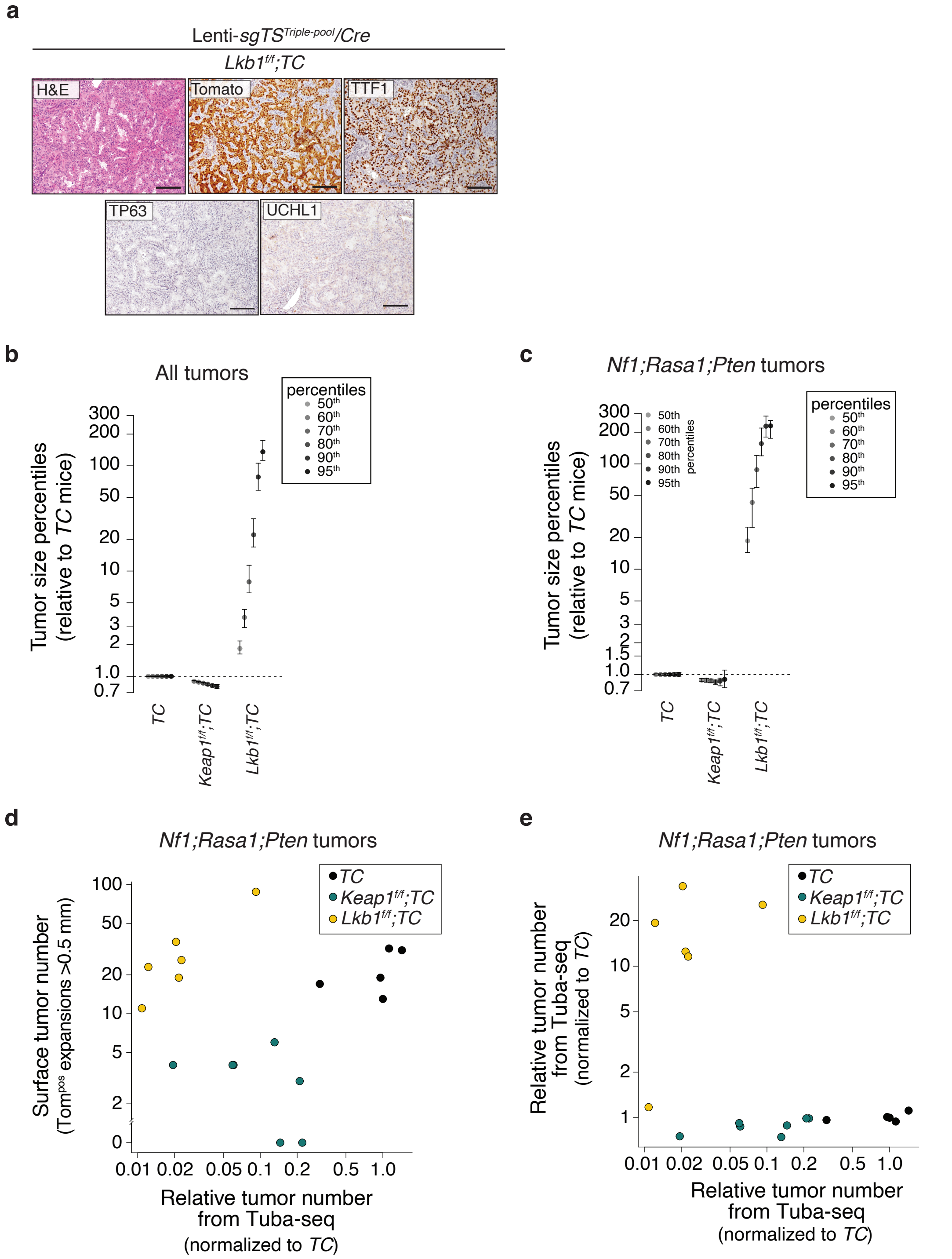


**Supplementary Figure 5. Inactivaton of the *Keap1* or *Lkb1* differentially impacts *Nf1;Rasa1;Pten* oncogene-negative tumor initiation/early expansion and tumor size**

**a,** Representative H&E and Tomato, TTF-1 (adenocarcioma marker), P63 (squamouse cell carcinoma marker), and UCHL1(small cell lung cancer marker) stained sections of a lung tumor from a *Lkb1*$^{flox/flox}$*;TC* mouse. Scale bar = 100 µm.

**b-c,** Tumor sizes at indicated percentiles relative to the same percentile of tumor sizes in TC mice. Whiskers show 95% confidence intervals from bootstrap resampling.

**d,** Comparison of Surface tumor number (from direct counting of Tomato$^{positive}$ tumors >0.5mm in diameter) with the Relative tumor number from Tuba-seq (the number of clonal exapansions with >50 neoplastic cells relative to the median of *TC* values). Each dot represents a mouse. Data from *Nf1;Rasa1;Pten* triple mutant tumors is shown.

**e,** Comparison of the Relative tumor size determined from Tuba-seq (the median number of neoplastic cells relative to the median of *TC* values) with the Relative tumor number from Tuba-seq (the number of clonal expansions with >50 neoplastic cells relative to the median of *TC* values). Each dot represents a mouse. Data from *Nf1;Rasa1;Pten* triple mutant tumors is shown.

**a**

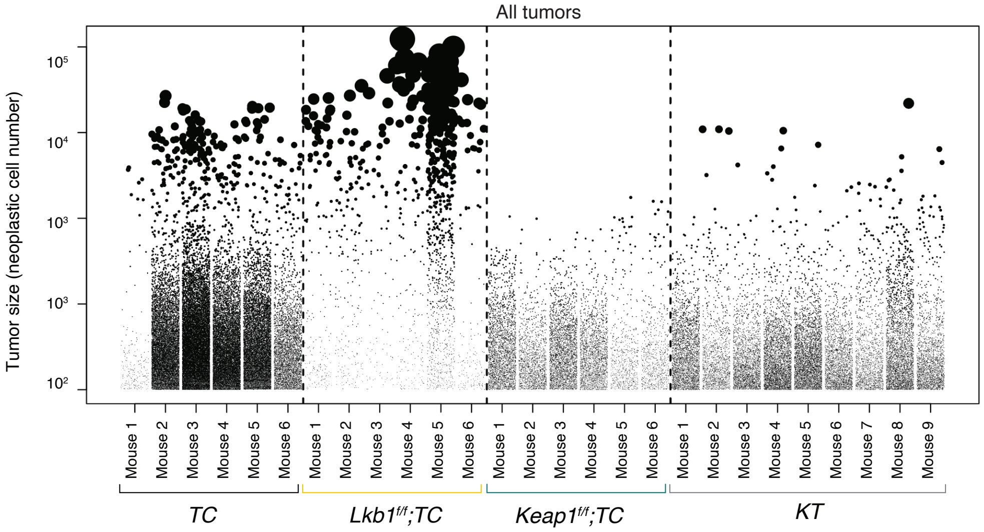


**b**

| | *TC* | *Lkb1$^{f/f}$;TC* | *Keap1$^{f/f}$;TC* | *KT* |
|---|---|---|---|---|
| *Nf1;Rasa1;Pten* | 2901 | 108 | 496 | 537 |
| *Nf1;Rasa1* | 669 | 39 | 90 | 131 |
| *Nf1;Pten* | 3198 | 66 | 526 | 594 |
| *Nf1* | 1974 | 21 | 315 | 697 |
| *Rasa1;Pten* | 1168 | 22 | 199 | 302 |
| *Rasa1* | 1854 | 24 | 284 | 552 |
| *Pten* | 825 | 12 | 137 | 223 |
| *Inert* | 1323 | 13 | 194 | 423 |

**Supplementary Figure 6. *Keap1* and *Lkb1* inactivation dramatically impacts the number and size of tumors initiated with Lenti-sgTS$^{triple\text{-}pool}$/Cre.**

**a,** Gitter plot of tumor sizes in each mouse of each genotype. Each dot represents a tumor (>50 neoplastic cells). Sizes of dots scale with tumor size. Each column represents a mouse. All tumors generated with any of the lentiviral vectors are shown.

**b,** Average total number of clonal tumors with each lentiviral vector in each mouse strain per mouse. Tumor numbers are the sum of all tumor with the given lentiviral vector avcross all mice of each genotype, divided by the number of mice of that genotype. Tumors are defined as clonal expansions estimated to have >50 neoplastic cells. These numbers reflect the number of barcoded tumors prior to removal of tumors to reduce the impact of multiple transduction events (see **Methods**).

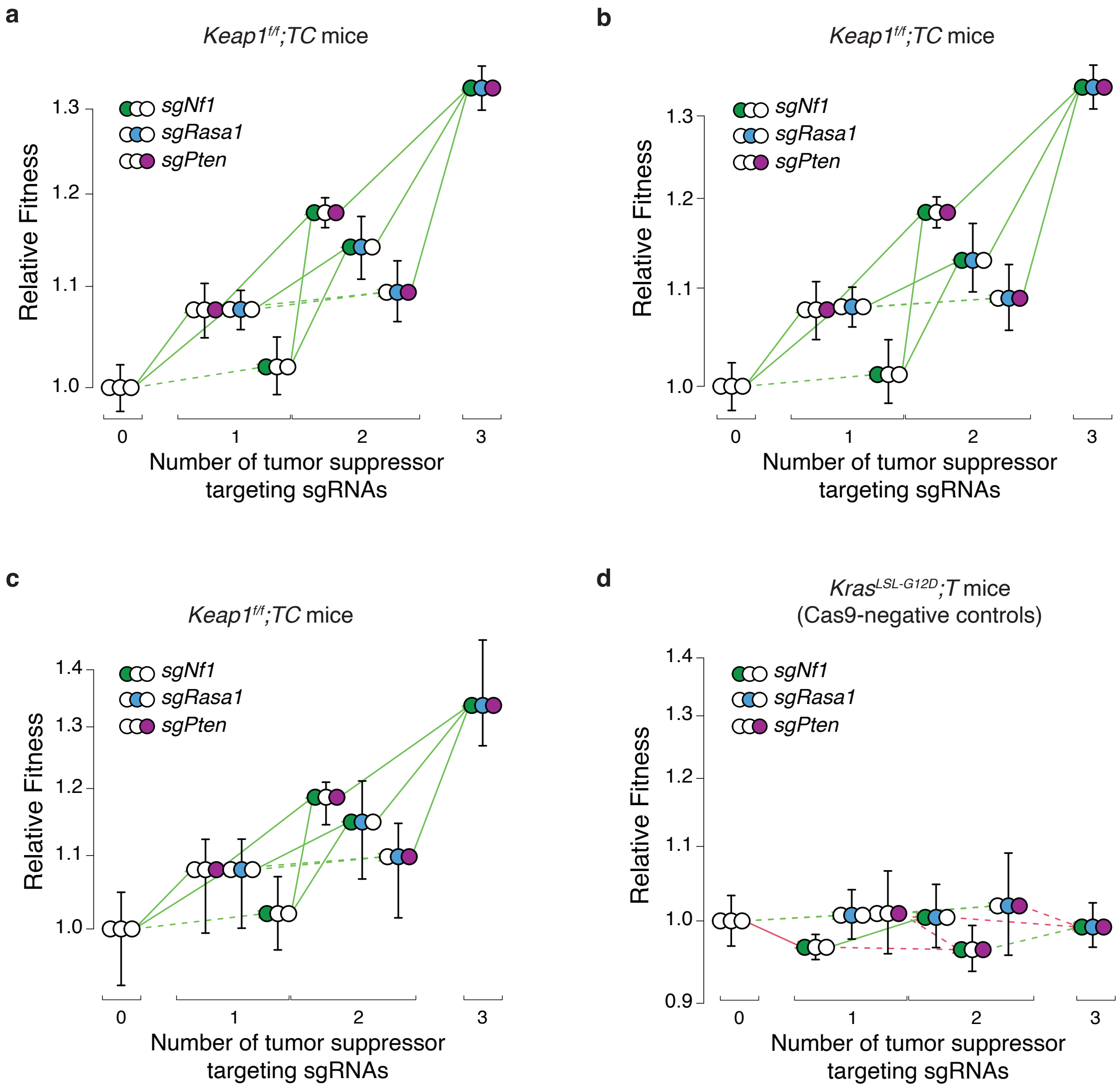


**Supplementary Figure 7. Estimates of fitness are robust to methods that incorporate different methods for multiple transduction correction as well as different methods of**

**a-c,** Fitness landscape of tumors in $Keap1^{f/f}$*;TC* mice without any multiple transduction corrrection (**a**), with multiple transduction using method #2 (Methods)(**b**) and with multiple transduction correction as in Figure 4b but with 95% confidence intervals calculated through bootstrap resampling of both mice and tumors (**c**). Green arrows indicate increased fitness, red arrows indicate reduced fitness, solid line indicates significance (p-value $< 0.05$). Whiskers in **b and c** show 95% confidence intervals from bootstrap resampling of tumors.

**d,** Fitness landscape for $Kras^{LSL\text{-}G12D}$*;T* (Cas9-negative control) mice. Fitness for tumors of each single and double mutant genotype, as well as those with all three tumor suppressor targeting sgRNAs are shown relative to the triple inert vector. Green arrows indicate increase in fitness, red arrows indicate reduced fitness, solid line indicates significance (p-value $< 0.05$). Whiskers show 95% confidence intervals from bootstrap resampling of tumors.

Fitness for tumors of each single and double mutant genotype as well as those with all three genes inactivated are shown relative to inert. Green arrows indicate increased fitness, red arrows inducate reduced fitness, solid line indicates significance (p-value $< 0.05$). Whiskers show 95% confidence intervals.
